# Digital twin reveals combinatorial code of non-linear computations in the mouse primary visual cortex

**DOI:** 10.1101/2022.02.10.479884

**Authors:** Ivan Ustyuzhaninov, Max F. Burg, Santiago A. Cadena, Jiakun Fu, Taliah Muhammad, Kayla Ponder, Emmanouil Froudarakis, Zhiwei Ding, Matthias Bethge, Andreas S. Tolias, Alexander S. Ecker

## Abstract

More than a dozen excitatory cell types have been identified in the mouse primary visual cortex (V1) based on transcriptomic, morphological and *in vitro* electrophysiological features. However, the functional landscape of excitatory neurons with respect to their responses to visual stimuli is currently unknown. Here, we combined large-scale two-photon imaging and deep learning neural predictive models to study the functional organization of mouse V1 using *digital twins.* Digital twins enable exhaustive *in silico* functional characterization providing a *bar code* summarizing the input-output function of each neuron. Clustering the bar codes revealed a continuum of function with around 30 modes. Each mode represented a group of neurons that exhibited a specific combination of stimulus selectivity and nonlinear response properties such as cross-orientation inhibition, size-contrast tuning and surround suppression. These non-linear properties were expressed independently spanning all possible combinations across the population. This combinatorial code provides the first large-scale, data-driven characterization of the functional organization of V1. This powerful approach based on digital twins is applicable to other brain areas and to complex non-linear systems beyond the brain.

## Introduction

Understanding the functional organization of the primary visual cortex (V1) has been a longstanding goal in neuroscience. It has long been known that V1 extracts information about local orientation Hubel & Wiesel (1959), often in a phase-invariant manner (Hubel & Wiesel, 1962). Researchers have described additional V1 nonlinearities, including direction selectivity (Adelson & Bergen, 1985) and various forms of nonlinear contextual modulation (Blakemore & Tobin, 1972; Cavanaugh et al., 2002; DeAngelis et al., 1992; Gilbert & Wiesel, 1990; Heeger, 1992; Lamme, 1995; Morrone et al., 1982). However, although we know many of the building blocks of V1 function, we do not know how they are organized at the population level.

First, we do not know whether there exists a distinct number of functional cell types, each of which implements a specific computation, or whether there is a continuum of function, where cells do not fall into discrete types. Second, independent of whether V1 functions are discrete or form a continuum, we currently do not know how the different nonlinear effects described previously are organized at the population level: are they strongly correlated – for instance because they are caused by a common computational mechanism – or are they present independently of each other within the population?

A major roadblock in revealing the functional organization of V1 has been that traditional experiments probing the well-known nonlinear mechanisms do not scale well. Large-scale population recordings are inefficient, because stimuli need to be optimized to an individual neuron’s receptive field location, preferred orientation and spatial frequency. In addition, probing all nonlinear mechanisms in the same neurons is difficult because only a limited number of stimuli can be shown in an experiment.

We have overcome these limitations by combining largescale population recordings with natural stimuli and training high-performance predictive models based on deep neural networks (Antolík et al., 2016; Batty et al., 2016; Cadena et al., 2019; Cotton et al., 2020; Klindt et al., 2017; Lurz et al., 2020; Sinz et al., 2018; Walker et al., 2019). These models are capable of jointly modeling thousands of neurons in a completely data-driven way providing a digital twin: an *in silico* approximation of the function of primaryvisual cortex (Fig. 1A). First, this approach allows us to quantify the similarity of neurons’ response properties on the set of natural stimuli by computing a compact, lowdimensional vector representation of each neuron’s function (its bar code). This representation is independent of the neuron’s receptive field location and its preferred orientation and provides an unbiased metric to measure the similarity of two neurons’ functions. It therefore provides a principled way to study the functional organization of V1. Second, the digital twin allows us to carry out experiments with arbitrary stimuli *in silico,* essentially without limitations of experimental time to generate hypothesis which can then verified back *in vivo* using the inception loop paradigm (Bashivan et al., 2019; Walker et al., 2019). This systematic functional analysis allows us to gain interpretable insights from the model and link to existing literature.

**Fig. 1.**
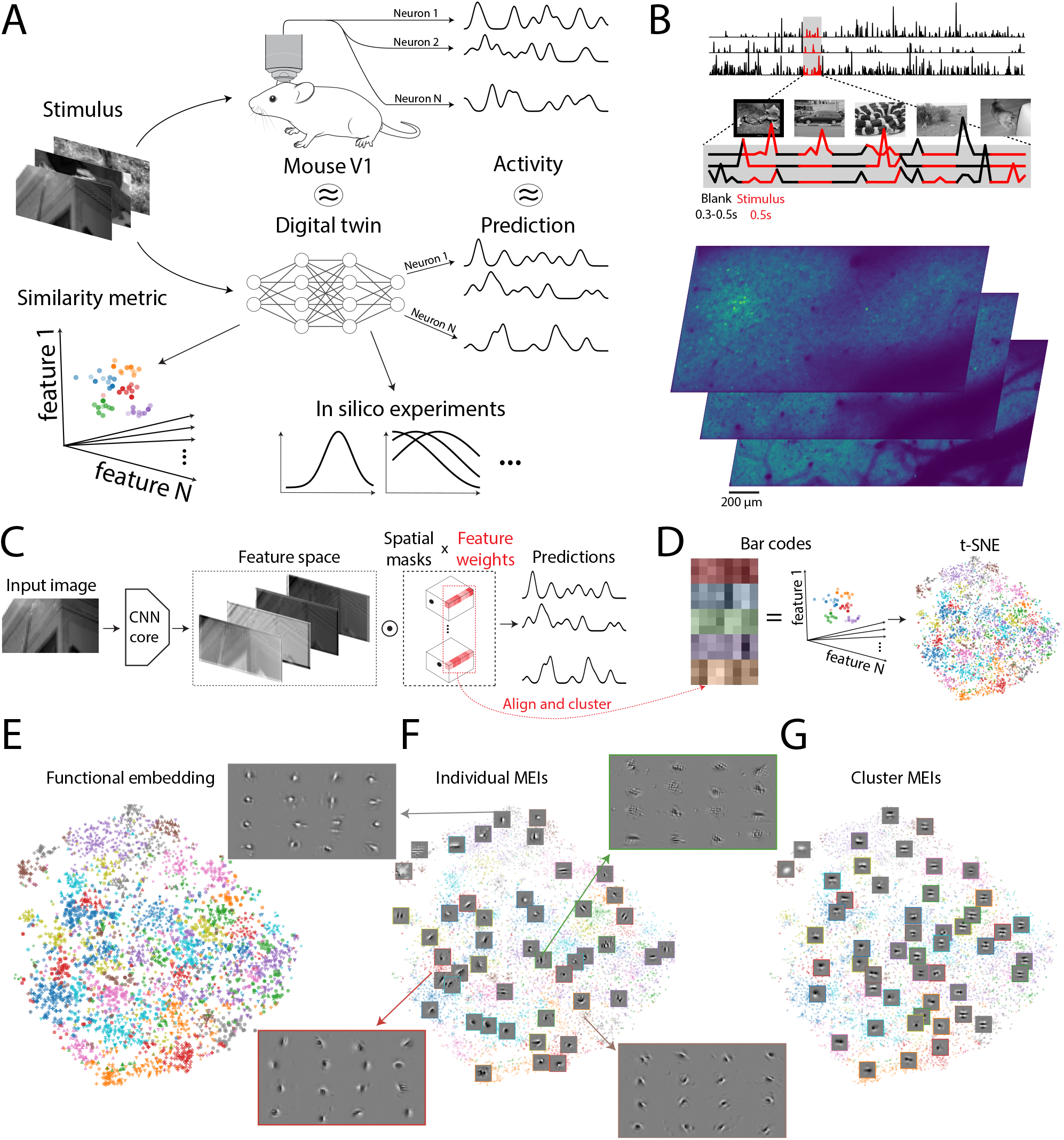
**A.** Overview of our method. We presented natural images to a mouse and recorded corresponding responses of a large population of neurons in the primary visual cortex. This dataset allowed us to build a “digital twin” model of the mouse primary visual cortex which provided a functional similarity metric between the neurons, as well as enabled us to perform *in silico* experiments. **B.** Data recording paradigm. We presented an alternating sequence of 4692 natural and blank images to seven different mice. We showed natural images for 0.5s and blank ones for a random duration between 0.3s and 0.5s. We presented an additional test set of 100 images 10 times each. We recorded responses of 5 to 8 thousand V1 L2/3 neurons depending on the scan using a wide-range two-photon microscope. Processed calcium traces for three randomly chosen neurons are shown in the top, and raw scans at three different depths are shown in the bottom. **C.** Model fitting paradigm. We pooled the data from all 7 mice in a single dataset and fitted a rotation-equivariant CNN model to predict the recorded neural activity. The model consists of a rotation-equivariant convolutional core shared across neurons and neuron-specific linear readouts. For each neuron the readout is decomposed into a spatial mask encoding the spatial location of its receptive field and a vector of feature weights encoding predictive CNN features for this neuron. Feature weights can be thought of as “bar codes” summarizing a neuron’s function. **D.** Functional clustering. We collected the feature weights for all neurons into a single matrix (row-wise) and aligned its rows by cycling shifts to remove the differences due to different preferred orientations of the neurons. We then clustered the rows of the aligned feature weights matrix into 50 clusters using the k-Means algorithm. The aligned feature weights and the clusters are visualized using a 2D t-SNE embedding. **E.** A 2D t-SNE functional emedding of the recorded neurons colored according to the cluster assignment. **F.** Examples of MEIs of 16 best predicted neurons in 4 different clusters alongside examples of MEIs of other neurons on top of a t-SNE embedding. **G.** Examples of cluster MEIs of other neurons on top of a t-SNE embedding.

We found that the functional organization is not entirely uniform, revealing a number of high-density modes. We therefore used the bar codes to cluster functionally similar neurons, which allowed us to analyze the neurons’ functional properties at the cluster level.

Crucially, our analysis revealed that classical non-linear properties of neurons in V1 are expressed independently of each other. For instance, knowing the extent of a neuron’s non-linearity along the simple-complex cell axis does not provide much information about its degree of surround suppression or cross-orientation inhibition. Moreover, there exist functional clusters expressing all combinations of nonlinear properties (including none or all), suggesting that V1 neurons might be described with a combinatorial code in the space of basic nonlinear computations.

Overall, our results suggest the following answers to the two questions posed above. The functional organization of V1 appears to form a continuum; however, it is not uniform and there are high-density modes in the space of V1 neurons’ functions. This organization is consistent with recent work using transcriptomic, morphological, and electrophysiolocal properties, which showed that cortical neurons are organized in families with a continuum of properties within them rather than distinct cell types Gouwens et al. (2020); Network (2021); Scala et al. (2021). With respect to classical nonlinearities, V1 neurons can be described by a combinatorial code where each nonlinear computation is expressed along an independent axis across the population. Such factorized codes have computational advantages such as higher coding capacity (Fusi et al., 2016).

## Results

### Large-scale recording and predictive modeling

We recorded the activity of more than 45,000 excitatory neurons in layer 2/3 of the primary visual cortex of seven mice using a wide-field two-photon microscope (Sofroniew et al., 2016, Fig. 1A). While we imaged, the mice were head-fixed on a linear treadmill and were viewing natural images, which covered roughly 120° × 90° of their visual field (Fahey et al., 2019; Walker et al., 2019). Next, we selected up to 2,000 neurons from each mouse and fitted a single predictive model for all mice (Lurz et al., 2020). The model is based on a convolutional neural network (CNN). It takes as input the image on the screen and outputs a prediction of the response of each neuron (Fig. 1C). The model achieved single trial test correlation of 0.42, and oracle correlation of 0.69. From this model, we obtained a 128-dimensional vector representation of each neuron’s function. These vectors can be thought of as “bar codes” summarizing the neuron’s stimulus-response function (Fig. 1 D).

To describe the neurons’ functional diversity, we removed two well-known factors of variation across V1 neurons: receptive field position and preferred orientation. The bar codes we obtained from our model were independent of receptive field position and preferred orientation of the neuron: if the responses of two neurons could be made identical by applying a constant shift and rotation to all images, these two neurons would obtain the same bar code. We achieved this property by using a rotation-equivariant CNN (Ecker et al., 2019; Ustyuzhaninov et al., 2020). Having bar codes that are independent of receptive field location and preferred orientation is extremely useful, because it removes two “trivial” axes of variation and allows us to focus on and visualize more subtle aspects of the neurons’ selectivity or nonlinear processing.

### Predictive modeling reveals functional clusters

We first asked whether V1 neurons are organized into discrete functional types or rather form a continuum. A 2D t-SNE embedding (van der Maaten & Hinton, 2008) of the bar codes (Fig. 1 E) revealed several modes – or regions of high density. These modes correspond to groups of functionally similar neurons. While there is no strong evidence for discrete functional types, it is not a uniform distribution either. We performed k-means clustering (MacQueen et al., 1967) (using 50 clusters) to identify the modes of the distribution and simplify downstream analysis.

### Neurons within functional clusters have similar MEIs

Given that V1 neurons can be organized into functional clusters, we aim at characterizing these clusters. We start by computing the preferred stimulus of each neuron, sometimes referred to as the most exciting image (MEI), which is optimized, using the model, to maximize the neuron’s predicted activity (Fig. 2A). They have been shown to provide a faithful snapshot of neural computations (Bashivan et al., 2019; Walker et al., 2019), and therefore provide convenient single-image visualizations of a neuron’s selectivity. Neurons within the same cluster had similar MEIs (up to location and rotation), while MEIs of neurons in different clusters tended to be different (Fig. 1F). This result provides a first piece of evidence that the clusters are a meaningful way of describing the functional organization of V1.

**Fig. 2.**
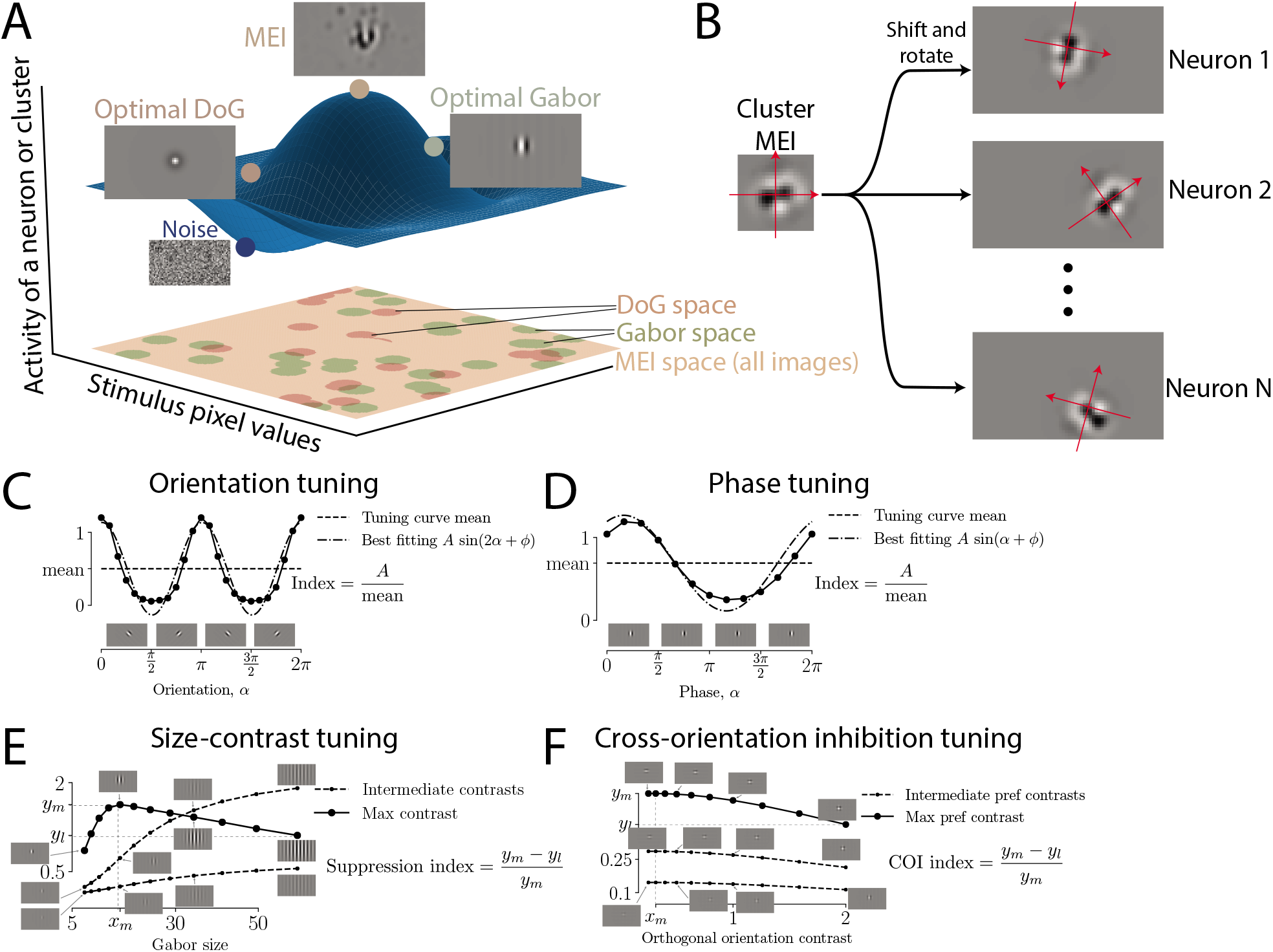
**A.** Optimal stimuli. We maximize activity of a neuron or average activity of a cluster with respect to an input stimulus which is constrained to belong to a certain image space. An illustration shows a response surface of a neuron of a cluster and maximum values on the this surface while restricting ourselves to a specific image space shown in the XY plane. The space of possible MEIs contains all images, while the spaces of possible Gabors and DoGs are subsets of all images. **B.** Average cluster activity for a given stimulus is computed by averaging the responses of neurons in the cluster to the stimuli shifted and rotated to match the location of the receptive field and preferred orientation of each neuron in that cluster. **C-D.** Examples of orientation and phase tuning curves for a single neuron. We vary orientation and phase of an optimal Gabor while keeping all other parameters fixed to generate stimuli for orientation and phase tuning experiments. The numerical tuning indices for these tuning curves are computed by fitting a sine curve and taking the ratio of its amplitude to the mean of the tuning curve. **E.** Example of a size-contrast tuning curve for a single neuron. The stimuli for the size-tuning experiment are constructed by varying size and contrast of an optimal Gabor while keeping all other parameters fixed. The suppression tuning strength is computed for a tuning curve corresponding to the highest contrast as a relative decrease of activity when increasing the size of the Gabor beyond the size corresponding to the maximum value of the curve. The contrast tuning strength is computed analogously by transposing size and contrast, i.e. by considering a tuning curve corresponding to the largest size as a function of the contrast. **F.** Example of a cross-orientation inhibition (COI) tuning curve for a single neuron. The stimuli for this experiment are called plaids and constructed by overlaying the optimal Gabor and the Gabor orthogonal to it in different contrasts. The COI tuning strength index is computed using a tuning curve corresponding to the highest preferred contrast analogously to the suppression index for the size-contrast experiment.

Note that our data does reproduce the large degree of heterogeneity of MEIs and their frequent striking deviations from Gabor filters (Fig. 1F) that have been reported previously for mouse V1 (Walker et al., 2019).

### Cluster MEIs visualize functional similarities between the neurons

To focus more on commonalities of functionally similar neurons, we next computed optimal stimuli at the cluster level. To do so, we computed cluster MEIs: image templates that maximize average activity of neurons in each cluster when accounting for each neuron’s receptive field location and preferred orientation (Fig. 2B). Cluster MEIs show a systematic variation along the different axes of the t-SNE embedding (Fig. 1G): Neighboring clusters tend to have visually similar cluster MEIs. There appears to be a global pattern in the t-SNE space with clusters on the right having oriented, Gabor-like MEIs with higher frequencies and multiple cycles within the envelope, while those in the middle having lower frequencies and fewer cycles and those towards the bottom left having more symmetric and circular MEIs.

Cluster MEIs visualize common patterns of cluster computations, and since they are designed to capture similarities rather than differences between the neurons, they exhibit less variability than individual MEIs. Quantitatively this amounts to cluster MEIs driving the neurons to around 60% of activity of their individual MEIs.

### In silico experiments provide an interpretable characterization of functional clusters

While MEIs provide convenient visualizations of a neuron’s or a cluster’s computations, they capture only a single point – the maximum – of the tuning function (Fig. 2A). Our predictive model, however, provides a prediction for arbitrary stimuli. We used the model as an *in silico* replica of V1 to perform experiments. Unlike with experiments in the real brain, in the model we are not limited in terms of experimental time. This allowed us to replicate a number of classical experiments *in silico* and compute tuning curves with respect to a variety of different non-linear properties. Specifically, we used Gabor stimuli whose parameters were optimized for each cluster to quantify strength of orientation selectivity (Fig. 2C), phase invariance (Fig. 2D), size-contrast tuning (Fig. 2E) and cross-orientation inhibition (Fig. 2F). Optimizing Gabors for clusters rather than individual neurons prevented the possibility of a few neurons within a cluster having very stimuli in comparison to the rest of the neurons in the cluster due to optimization instability. Furthermore, the cluster optimal Gabors drive the neurons to 87% of their individual optimal Gabor activity thus these stimuli are only mildly suboptimal for individual neurons.

Size-contrast tuning curves reveal non-linear surround suppression effects (Born & Tootell, 1991; DeAngelis et al., 1992), while cross-orientation inhibition is a nonlinear interaction that arises when two orthogonal Gabor patterns are superimposed (Morrone et al., 1982). In addition, we computed the optimal center-surround stimulus (difference of Gaussian; Fig. 2A) and quantified the degree of response nonlinearity using a generalized linear model baseline (see methods). From these in silico experiments, we obtain a sample from the joint tuning distribution for more than 10,000 neurons. This enables us to study the statistical dependencies between the different nonlinear effects, overcoming the limitations of previous *in vivo* experiments that could only study each effect in isolation.

### In silico experiments reveal shared tuning properties within functional clusters

The results from the set of in silico experiments support the functional clustering (Fig. 3). Many of the modes in the t-SNE embedding are distinguishable based on one or more of the tuning properties. In contrast, neurons *within* most of the clusters exhibit similar tuning strengths to the different types of non-linarities: all 50 clusters were significantly different from the overall population tuning distribution based on at least one tuning property in Fig.3 and 15 clusters were significantly different based on all properties (twosided Kolmogorov-Smirnov test at *α* = 0.01; in the case of random cluster assignments these values are 0 and 1 respectively).

**Fig. 3.**
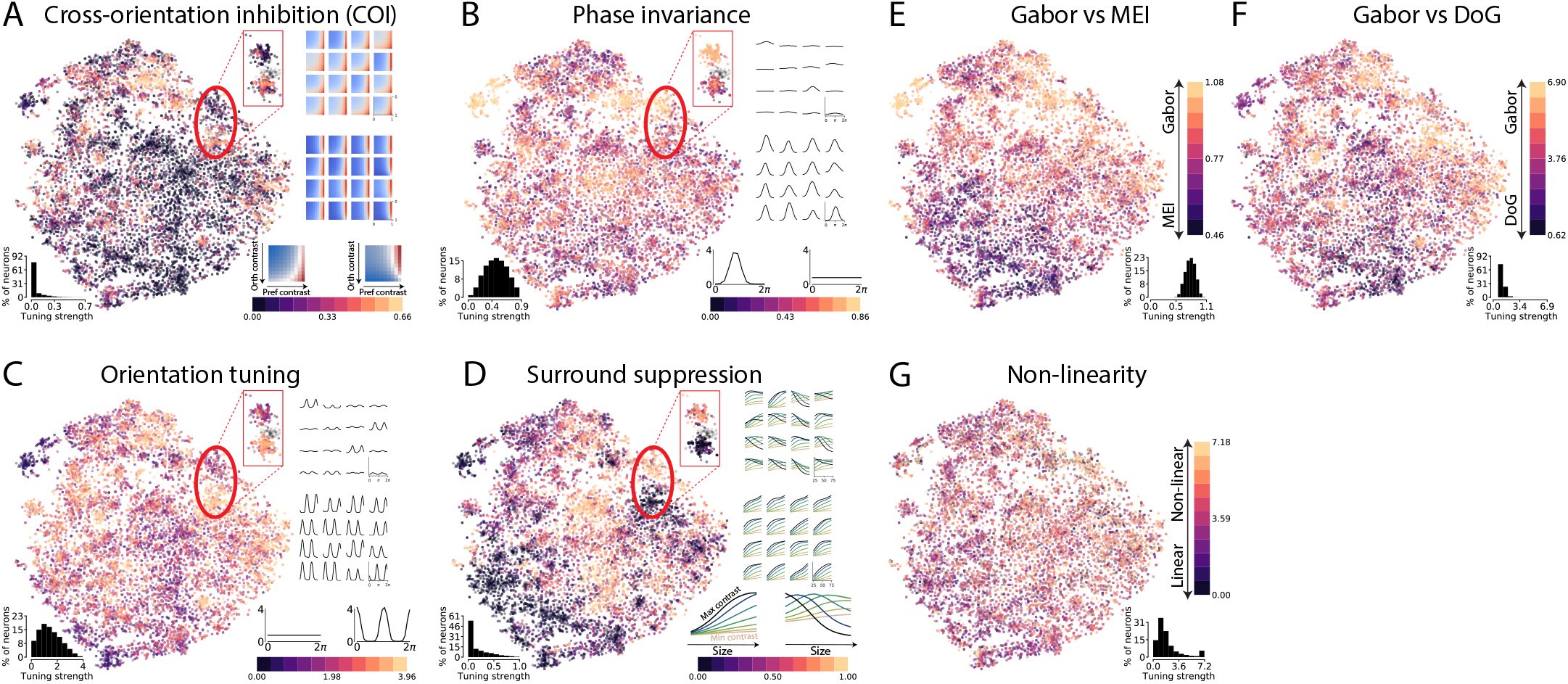
Results of the in silico experiments. **A-D:** The scatter plots show t-SNE embeddings colored according to the tuning strengths of cross-orientation inhibition, phase invariance, orientation tuning and surround suppression experiments. In the bottom left corners of t-SNE embeddings we show histograms of distributions of the tuning strengths. We additionally show the tuning curves for the 16 best predicted neurons in two different clusters in a separate close-up panel, as well as examples of tuning curves of high and low tuning strengths alongside the colorbar. **E-G:** t-SNE embeddings colored according to the tuning strength of Gabor vs MEI, Gabor vs DoG and non-linearity experiments along with histograms of tuning strengths.

### Non-linear tuning properties are independent of each other

Next, we investigated how different non-linear properties relate to each other. Qualitatively, it appears that different non-linear properties have different distributions (Fig. 3; compare the color patterns in the t-SNE plots). This qualitative impression is also confirmed by quantitative metrics (Fig. 4A). As expected, non-linearity is correlated with specific nonlinear properties like phase invariance, surround suppression and cross-orientation inhibition, but not with orientation selectivity. In addition, orientation selectivity is correlated with phase invariance and cross-orientation inhibition. Importantly, there is no correlation between the three non-linear properties phase invariance, surround suppression and cross-orientation inhibition, suggesting that these properties are independently exhibited from each other. This result is surprising, since cross-orientation inhibition and surround suppression have both been hypothesized to arise from a common mechanism: divisive normalization (Carandini & Heeger, 2012).

**Fig. 4.**
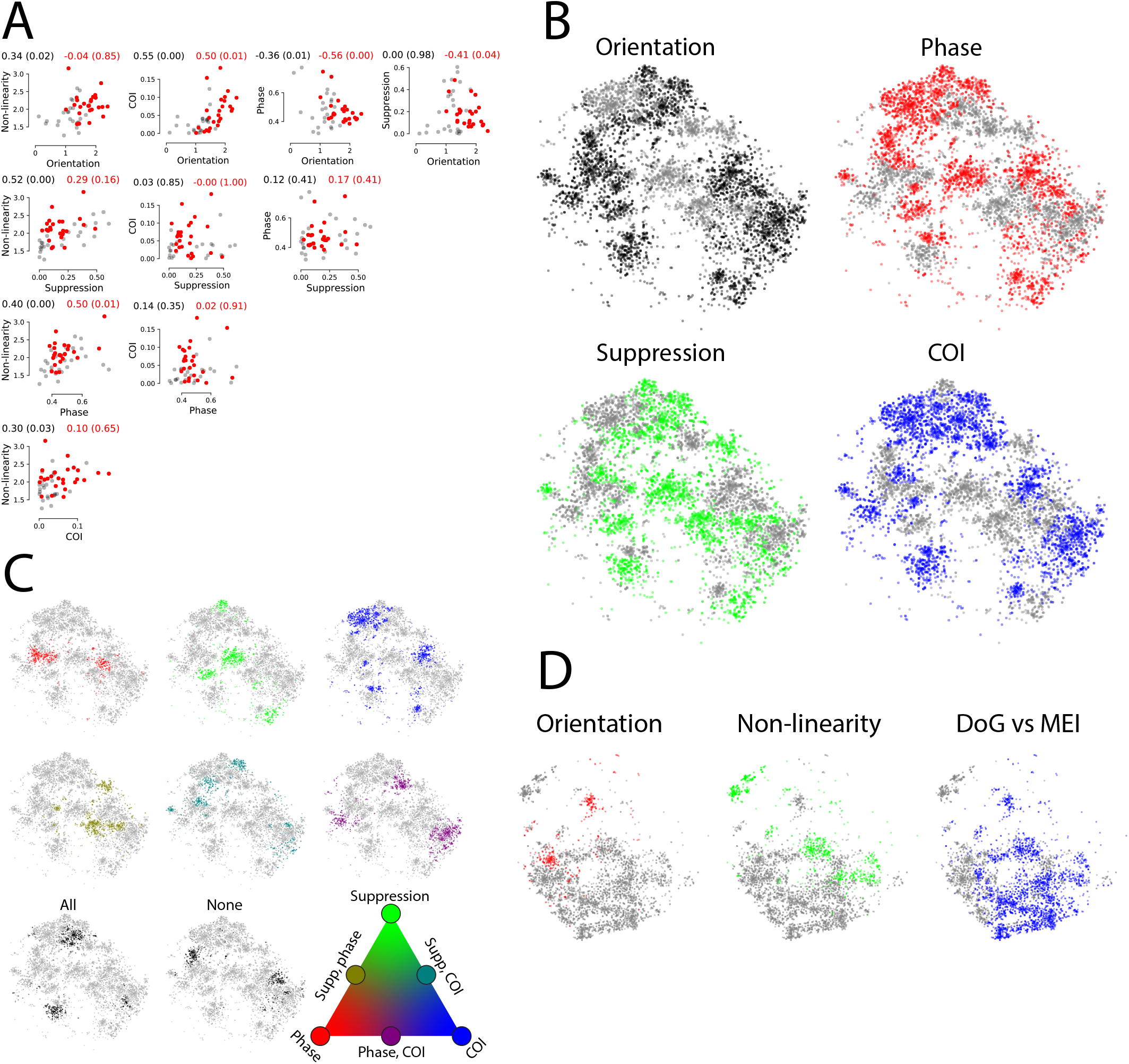
**A.** Pairwise distributions ofthe in silico tuning properties. Dots in each of the plots correspond to clusters (50 dots in total) showing average tuning strength of neurons in the cluster. Red dots correspond to clusters with the above average value of the Gabor vs MEI tuning strength (i.e. those clusters for which optimal Gabors are good stimuli relative to MEIs; in the following called Gabor-like clusters), gray dots correspond to all other clusters. Correlation coefficients and the p-values (in brackets) under the null hypothesis that the correlation is zero are shown above the plots for Gabor-like clusters (red) and for all clusters (black). **B.** Subsets of t-SNE embeddings corresponding to Gabor-like clusters colored according to the binarized strengths of orientation, phase, suppression and plaids tuning. Grey clusters correspond to low tuning strength (average cluster tuning strength is less than entire population tuning strength average value), clusters of the other color correspond to high tuning strength. **C.** Subsets of t-SNE embeddings corresponding to Gabor-like clusters showing 8 possible combinations of low/high values of phase, suppression and plaids tuning. The color code is illustrated with a color triangle; clusters colored with the colors in the vertices of the triangle exhibit high value of the corresponding tuning property and low values of other two properties. Clusters colored with the colors in the edges of the triangle exhibit high values of the tuning properties in the adjacent vertices and low value of the other tuning property. **D.** Subsets of t-SNE embeddings corresponding to non Gabor-like clusters colored according to the binarized strengths of orientation, non-linearity and DoG vs MEI tuning.

Because phase invariance, cross-orientation inhibition and surround suppression are tested using oriented Gabors as stimuli, we restricted our subsequent analyses to those clusters for which optimal Gabors are decent stimuli in comparison to MEIs (i.e. clusters with above average values of the Gabor vs MEI index). We binarized the tuning indices for these non-linear properties assigning each cluster to either high or low tuning category (Fig. 4B) by setting a threshold at the average population tuning strength and examined the combinations of these three tuning properties (Fig. 4C). We can see there are clusters exhibiting every possible combination of the three binary tuning properties, suggesting that these properties indeed appear to be independent from each other, and that V1 neurons might employ a combinatorial code with respect to these nonlinearities.

For the analysis of remaining clusters (i.e. clusters with below average values of the Gabor vs MEI index, Fig. 4D) we considered orientation, non-linearity, and DoG vs MEI tuning. These clusters are mostly linear tuned and unoriented with their cluster MEIs ranging from mostly center-surround shapes in the bottom-right and central parts of the t-SNE space to various complex shapes in the left part of the t-SNE space.

### In silico tuning curves are good approximations of in vivo tuning

Since our analysis is based on in silico tuning curves, one concern could be that although the CNN model predicts neural activities with high accuracy, the corresponding in silico tuning curves might be different from the in vivo tuning curves we are aiming to approximate. We verified that it is not the case by directly comparing the in silico and in vivo tuning curves for the same neurons. Specifically, we recorded a dataset containing two V1 scans of the same neurons in the same mouse. We used natural images as stimuli for the first scan which allowed us to fit a rotation-equivariant CNN model. For the second scan we used Gabor stimuli allowing us to compute size tuning curves in vivo and compare them to the in silico tuning curves obtained from the model fitted to the first scan (Fig. 5A).

**Fig. 5.**
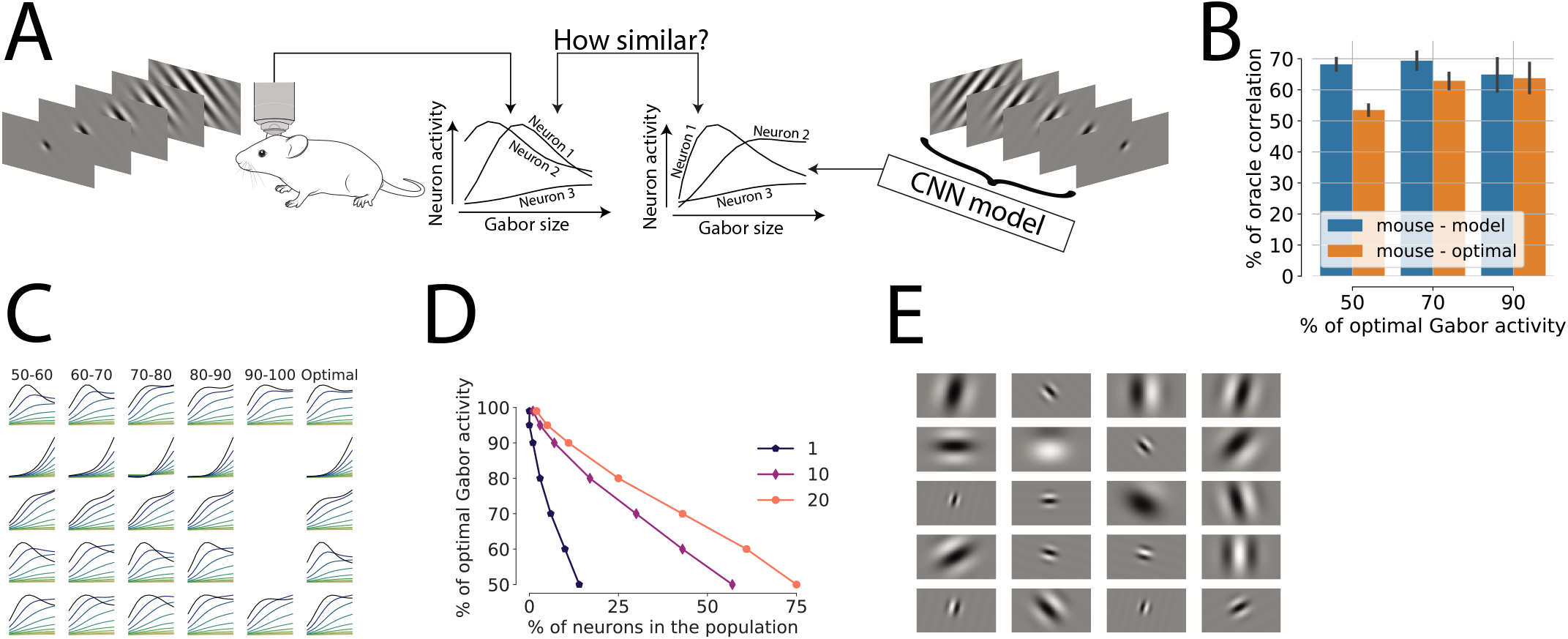
**A.** In vivo verification experiment paradigm. We compare in vivo Gabor size tuning curves to their in silico counterparts computed for the same population of neurons. **B.** Correlations between in vivo and in silico Gabor size tuning curves measured as percentage of oracle correlation (error bars shows the standard deviation). We measure correlation between two types of tuning curves. The “mouse – model” one corresponds to the in vivo and in silico tuning curves obtained using the same experimental stimuli, which are based on suboptimal Gabors (see main text for more details). In the “mouse – optimal” case, in vivo curves are computed using suboptimal Gabors, but the in silico curves are computed using optimal Gabors for the corresponding neurons. Moreover, these correlations are computed for different subsets of neurons (x-axis) chosen such that there exists an experimental stimulus for every neruon in the subset driving this neuron to a certain percentage of the optimal Gabor activity. **C.** Examples of Gabor size-contrast in silico tuning curves for 5 randomly chosen neruons (rows) computed using suboptimal Gabors activating the neuron to a certain percentage of the optimal Gabor activity (first 5 columns; numbers above the first row show the suboptimal Gabor activity as percentage of the optimal Gabor activity) and using the optimal Gabors (last column). **D.** Percentages of neurons in the population (x-axis) that can be driven to a certain percentage of the optimal Gabor activity (y-axis) using at least one of the Gabors in the sets of 1, 10 and 20 Gabors (different curves) chosen to maximize the number of neurons activated by these stimuli. **E.** 20 Gabor stimuli corresponding the 20 Gabors curve in panel **D** and used to construct size-tuning stimuli for the in vivo experiment.

The stimuli for the size tuning experiment should ideally be constructed based on optimal Gabors for every neuron, however, that is infeasible in vivo for a sufficiently large population of neurons. To overcome this limitation, we investigated to what extent optimal Gabors could be replaced with suboptimal ones. We found that tuning curves obtained using Gabors driving neurons to at least 50% of their optimal Gabor activities are very similar to the tuning curves obtained with optimal Gabors (Fig. 5C), and that only 20 different Gabors can be chosen to drive 75% of the population to at least 50% of optimal Gabor activities (Fig. 5D-E). This observation allowed us to use these 20 different Gabors as stimuli for the Gabor scan, which revealed that the correlation between in vivo and in silico tuning curves is about 70% of the oracle correlation (Fig. 5B), which is similar to the performance of the CNN model trained and tested on natural images.

## Discussion

We built a functional description of the mouse primary visual cortex based on neural representations in a high performing CNN model predicting responses of a large population of neurons on arbitrary natural images. Such an approach allows us to account for all aspects of the neuronal stimulus-response function captured by the model, instead of only a few nonlinear effects as in classical in vivo experiments with parametric stimuli. Thus, our analysis is not constrained by specific hypotheses of neural functions or the choice of parametric stimuli. An important limitation of our study is that we focus on the bottom-up aspects of stimulus processing; we did not consider how top-down, behavioral modulation of neural responses affects the responses of different neurons.

Our examination revealed that the V1 functional landscape can be described by around 30 modes of functionally similar neurons which, however, do not appear to be discrete cell types but rather high density areas in the continuous functional space. This finding is in agreement with various recent studies. For example, Scala et al. (2021) studied mouse primary motor cortex neurons based on transcriptomic and morpho-electric properties. They found that this brain area is organized into a few broad transcriptomic families with continuum of morpho-electric features in each family, making the authors question the existence of discrete transcriptomic cell types. Gouwens et al. (2020) conducted a similar study of interneurons in mouse primary visual cortex discovering both discrete and continuous variation of morpho-electric properties within the transcriptomic types. Overall, a growing body of literature suggests that mouse neocortex is organized in a complex way and cannot be adequately described by either discrete cell types or a uniform continuum of neurons. This notion of mouse neocortex organization is qualitatively different from the mouse retina, which exhibits discrete cell types (Baden et al., 2016), raising interesting directions for future research: how are other areas of neocortex organized based on various neural properties? and how do they relate to lower level brain areas projecting into the corresponding neocortical areas?

As we investigated the variability of individual neurons within the functional modes, we found visual differences between cluster and single neuron MEIs. We believe this is an expected consequence of clustering, which abstracts away individual neurons’ differences and reduces the complexity of the V1 functional space by focusing on similarities between neurons. To better understand the properties of functional modes, their correspondence to cluster MEIs and relate them to previous work, we performed in silico experiments, which revealed that neurons belonging to the same mode exhibit common response patterns. This observation suggests that neurons in the same functional cluster may form computational cliques important for downstream processing, offering an interesting direction for future research. For example, integrated approaches combining functional characterization using digital twins and connectomics data can determine the connectivity of neurons within and across functional clusters and commonalties of their inputs and projection targets (Bae et al., 2021).

Large-scale in-silico experiments allowed us to study statistical dependencies between various nonlinear phenomena known from single-neuron in-vivo experiments. We found these effects to be independent of each other, suggesting that V1 might employ a combinatorial code between modes of functionally similar cells. The mechanisms leading to such a code in V1 and their implications for downstream processing remain unclear. A speculation that lies at hand is that there might be a basis of independent non-linear computations serving different purposes in downstream processing, thereby building a foundation for specializations in higher visual areas. As a potential verification of this hypothesis and as a question in itself, future experimental work could investigate if neurons of the same functional cluster project to the same downstream area.

Finally, we verified that in silico experiments in a high-performing CNN model provide a good approximation of the in vivo tuning curves, thus substantiating our results based on the analysis of in silico tuning curves. Our in vivo verification is consistent with the findings of Walker et al. (2019) who report that in silico model MEIs also highly activate actual neurons. Overall, these observations suggest that high-perfroming CNN models can be considered “digital twins” of real neurons, a paradigm that has started being explored relatively recently but already provided significant insights into the brain and has a potential of becoming the main tool for future research.

## ACKNOWLEDGEMENTS

The research was supported by the German Federal Ministry of Education and Research (BMBF) via the Competence Center for Machine Learning (FKZ 01IS18039A); the German Research Foundation (DFG) grant EC 479/1-1 (A.S.E.), the Collaborative Research Center (SFB 1233, Robust Vision) and the Cluster of Excellence “Machine Learning – New Perspectives for Science” (EXC 2064/1, project number 390727645); the Bernstein Center for Computational Neuroscience (FKZ 01GQ1002); the National Eye Institute of the National Institutes of Health under Award Numbers U19MH114830 (A.S.T.) R01MH109556 (AST), P30EY002520, and the Intelligence Advanced Research Projects Activity (IARPA) via Department of Interior/Interior Business Center (DoI/IBC) contract number D16PC00003. The U.S. Government is authorized to reproduce and distribute reprints for Governmental purposes notwithstanding any copyright annotation thereon. Disclaimer: The views and conclusions contained herein are those of the authors and should not be interpreted as necessarily representing the official policies or endorsements, either expressed or implied, of IARPA, DoI/IBC, or the U.S. Government. The funders had no role in study design, data collection and analysis, decision to publish, or preparation of the manuscript.

## AUTHOR CONTRIBUTIONS

I.U., M.B., A.S.T and A.S.E. designed the study; J.F., T.M., K.P., E.F. and Z.D. performed imaging experiments and pre-processing of raw data; I.U., A.S.E. developed the in silico analysis framework; I.U., M.F.B., S.A.C. and A.S.E analyzed the data; I.U., M.F.B. and S.A.C. wrote the original draft; I.U., M.F.B, S.A.C. and A.S.E. reviewed and edited the manuscript with the input from M.B. and A.S.T.

## Methods

### Visual stimuli

We used gray-scale ImageNet images (Russakovsky et al., 2015) as visual stimuli in the data collection experiment. The number of images varied across the scans (Tab. 1) with 4692 images in the intersection, i.e. presented to a mouse in each of the scans. For the test set we used 100 images each repeated 10 times; for some scans and some images there were fewer repeats, in which case we resampled the recording to have 10 repeats in every scan. The screen was 55 × 31 cm at a distance of 15 cm, covering roughly 120° × 90°. Each image was presented for 500 ms followed by a blank screen lasting between 300 ms and 500 ms. The response of a neuron to a given stimulus is represented as a number of spikes in the time interval between 50 ms to 350 ms following the stimulus onset.

### CNN model and rotation-invariant clustering

We used the same architecture and training of the rotation-equivariant CNN model as in Ustyuzhaninov et al. (2020). For the clustering of aligned feature vectors (rotation-invariant clustering) we used the k-Means algorithm (MacQueen et al., 1967) with 50 clusters, which we empirically found to provide a good balance between clusters being small enough to contain similar neurons and the total number of clusters being relatively small.

### GLM model

We also fitted a GLM model (McCullagh & Nelder, 2019) to every neuron in the recorded dataset to evaluate the non-linearity of neurons or clusters (as measured by the non-linearity index, see below). We used Poisson likelihood as the noise model and log link function to ensure the predicted neural activities are non-negative. We cross-validated the *L*_2_ regularization coefficient for every neuron separately by considering 48 log-spaced values in [0.1,5].

### Optimal stimuli

Classical experiments that we aim at replicating in silico measure changes in neural activity in response to a certain type of transformations of an input stimulus (e.g. an orientation tuning experiment might use Gabor stimuli in different orientations). Ideally the input stimuli should be optimized for every neuron separately which is infeasible in vivo, but can be achieved in silico using a CNN model. In this section we describe the stimuli we use in the in silico experiments and how we optimize them for individual neurons or clusters.

#### Per neuron optimal Gabors

For every neuron we compute a Gabor stimulus which maximizes the predicted activity of this neuron. We parametrize such stimuli in terms of a spatial location (*r^x^,r^y^*), size *σ,* spatial frequency *v,* contrast *a,* orientation *φ* and phase *τ*. Specifically the value of the pixel in the *i*-th row and *j*-th column of an input image is defined using the following expression

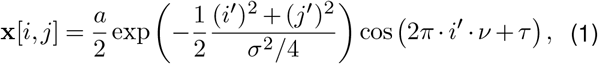

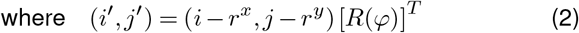

with *R*(*φ*) being a 2D rotation matrix by an angle *φ*.

For each neuron we iterate over a large set of Gabor stimuli parameterized according to (2) and record the parameter set corresponding to a stimulus with the highest CNN predicted activity. This process is illustrated in Fig. 2A.

#### Per cluster optimal Gabors

In addition to optimising Gabors for each neuron separately, we also find optimal Gabors for the entire clusters. The idea is to find a single Gabor stimulus that maximizes the average activity of neurons in the cluster and hence servres as a single image representations of a cluster computation. However, since we the clusters are explicitly constructed to contain neurons with different receptive field locations and preferred orientations, we constrain the stimuli to be identical for all neurons in the cluster apart from having neuron-specific spatial locations (i.e. receptive field centers) and orientations (see an illustration in Fig. 2A).

To find such an optimal Gabor for a specific cluster, we iterate over a large set of Gabors with locations and orientations set to a fixed value and all other parameters varying, generate stimuli for individual neurons by shifting and rotating the Gabor at the current iteration (Fig. 2A), and compute the average predicted activity of the neurons in the cluster. We call the Gabor stimulus (or rather the set of stimuli up to a spatial location and orientation) corresponding to the maximal average predicted cluster activity the cluster optimal Gabor.

#### Per neuron MEIs

An MEI is an input stimulus that activates the neuron the most, and as such serves as a useful visualization of the computation implemented by the neuron. It is important to keep in mind that an MEI is only a local maximum of a function mapping the input stimuli to the neural activity. While we aim at describing the entire function rather than only its maximum, MEIs nevertheless provide convenient and insightful summaries of computations performed by each of the neurons.

We compute MEI for every neuron separately by stating with a noise stimulus and iteratively optimising it to maximise the predicted activity of the neuron. Specifically, if the CNN prediction of activity of the *n*-th neuron when presented with a stimulus **x** is *f_n_*(**x**), we compute an MEI as a solution of the following optimisation problem:

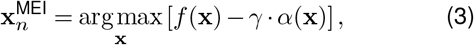

where *α*(**x**) is a regularisation function (e.g. *α*(**x**) = ||**x**||^2^) enforcing smoothness of the resulting MEI.

**Fig. 6.**
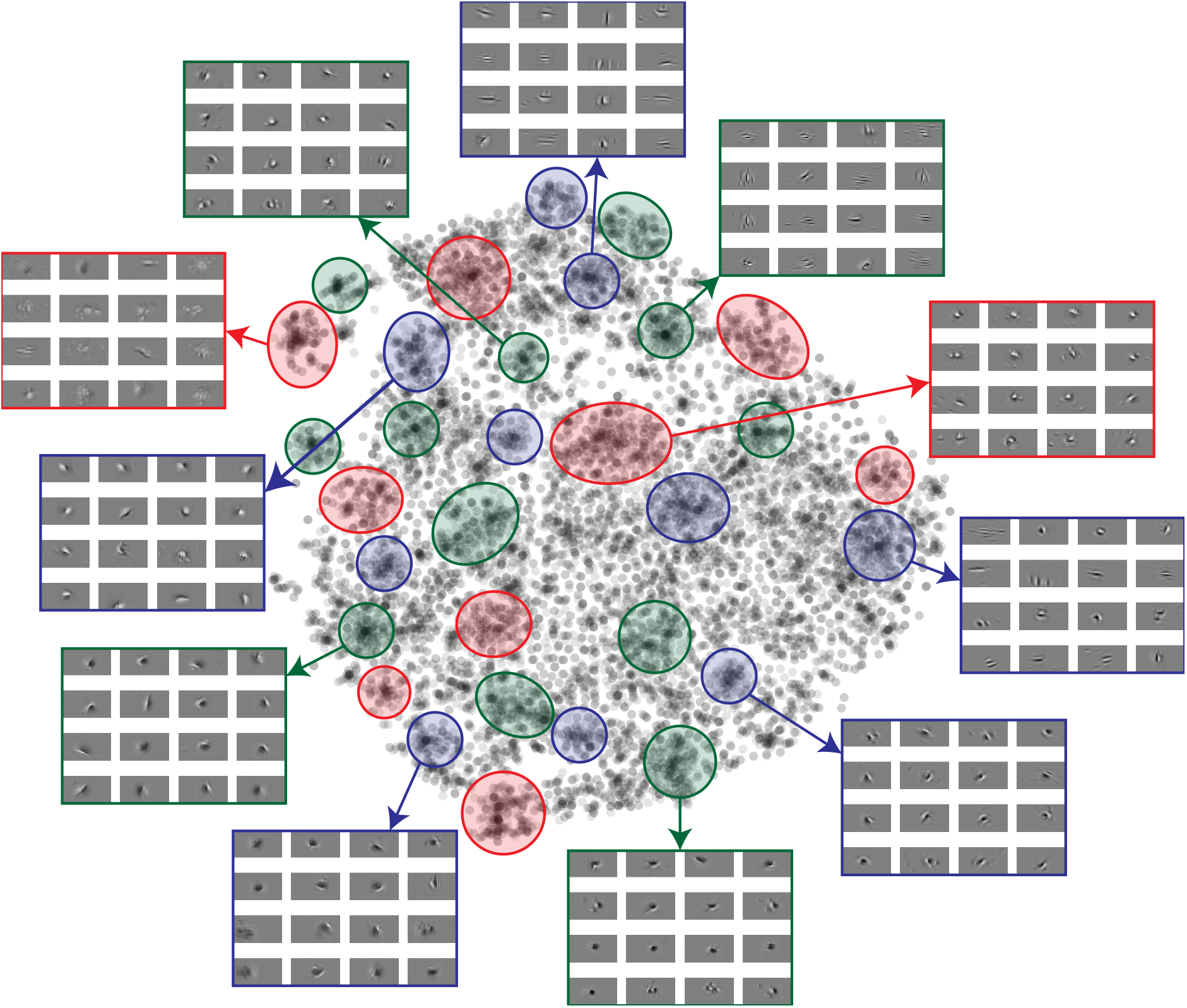
Functional modes in the t-SNE space.

**Table 1.** Summary of the individual mouse scans comprising the dataset used for the cell types analysis.

| Animal | # of neurons in the scan | Sampled neurons | # of train images | # of test images |
| --- | --- | --- | --- | --- |
| 1 | 5043 | 1047 | 5998 | 1999 |
| 2 | 5984 | 2000 | 5973 | 1090 |
| 3 | 7312 | 2000 | 4926 | 985 |
| 4 | 5335 | 2000 | 4994 | 999 |
| 5 | 8367 | 2000 | 4818 | 964 |
| 6 | 6045 | 2000 | 4982 | 995 |
| 7 | 8316 | 2000 | 4991 | 997 |
| Total |  | 13047 | 4692 | 1000 |

#### Per cluster MEIs

Such MEIs are computed jointly for all neurons in the entire cluster to maximise the average predicted activity of all neurons in the cluster. Similarly to cluster optimal Gabors, these stimuli are constrained to be identical for all neurons in the cluster apart from having neuron-specific spatial locations (i.e. receptive field centers) and orientations (Fig. 2A). To implement these constraints in practice, we decompose cluster MEIs in a steerable basis and iteratively optimise the coefficients in this basis to maximize the predicted average activity of neurons in the cluster. Therefore we use a parametric model of cluster MEIs rather than a non-parametric representation of single-neuron MEIs, which reduces the spaces of possible stimuli (for single-neurons MEIs this space consists of all possible images, while in this case it consists only of the span of the steerable basis), but allows us to compute stimuli rotations as linear transformations of the coefficients.

We consider the following parametrization of the stimuli

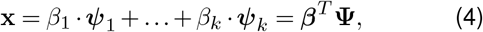

where **Ψ** = (*ψ*_1_,...,*ψ_k_*)^*T*^ is a vector of the first *k* basis functions in some steerable basis (we use Hermite polynomials). Rotations of images in such a parametrization correspond to linear transformations of coefficients *β*, specifically, a stimulus rotated by an angle *φ* can be written as

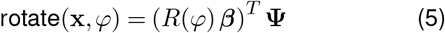

for a corresponding rotation matrix *R*(*φ*).

We denote a translation of an image **x** by *r^x^* pixels along the first axis and by *r^y^* pixels along the second axis as shift(**x**,*r^x^,r^y^*). Using center locations of optimal Gabors 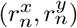 as receptive fields centers for corresponding neurons, we compute the cluster MEI 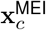 for a cluster *c* containing *m* neurons with indices *c*_1_,...,*c_m_* as a solution of the following problem

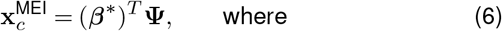

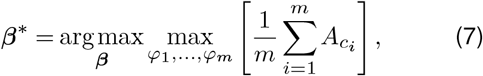

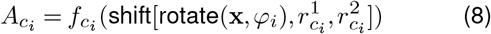

Examples of cluster MEI are shown in Fig. 3.

#### Optimal differences of Gaussians (DoG)

Another class of stimuli we consider are differences of Gaussians, which allow us to probe to what extent the receptive of a neuron has a center-surround structure. We parametrize such stimuli in terms of their spatial locations **r** = (*r*_1_;*r*_2_), sizes *σ*_cen_ and *σ*_sur_ of the center and surround Gaussians, as well as their relative contrasts *a*_cen_ and *a*_sur_ according to the following equation:

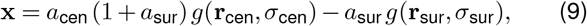

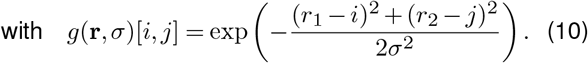

Similarly to optimal Gabors and MEIs, we find optimal DoG stimuli both for every neuron and cluster by iterating over a large set of parameter values and recording the parameter combination corresponding to the highest predicted activity of an individual neuron or average activity of all neurons in the cluster. In the case of per cluster stimuli, we constrain every neuron in the cluster to have exactly the same optimal stimulus apart from differences in spatial locations (Fig. 3A).

### In silico experiments

In this section we describe specific in silico experiments that we perform using the optimal stimuli discussed in the previous section.

#### Orientation and phase tuning

Optimal Gabors enable us to compute standard orientation and phase tuning curves. We use both per neuron and per cluster optimal Gabors to obtain stimuli for this experiment by varying the orientation and phase parameters and keeping all other parameters fixed. Examples of such stimuli are shown in Fig. 2B.

For every neuron we compute numerical indices reflecting the tuning strength of the neuron. Specifically, we compute the *F*_1_/*F*_0_ summary statistics for phase tuning and *F*_2_/*F*_0_ statistics. These statistics are ratios of the absolute values of the first and second (reflecting periods of 2*π* for phase tuning and *π* for orientation tuning) Fourier coefficients to the mean value of the tuning curve, or alternatively the ratios of amplitudes of the est fitting sine curves to the means of the tuning curves as illustrated in Fig. 2B. If **r**(**s**) = (*r*_1_(*s*_1_),...,*r_m_*(*s_m_*)) are responses of a neuron to Gabors with a parameter of interest (orientation of phase) taking values **s** = (*s*_1_,..., *s_m_*), these indices are defined as

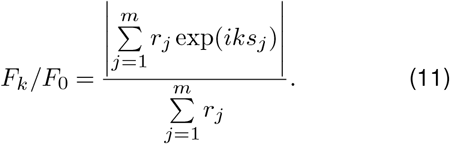

#### Gabor size-contrast tuning curves

We construct stimuli for this experiment by using per neuron or per cluster optimal Gabors with all parameters fixed except for the size and contrast as illustrated in Fig. 2B. The resulting size-contrast tuning curves allow us to characterize neural computations in terms of surround or contrast suppression effects, which have been widely studied in the existing literature.

Denoting predicted activity of *n*-th neuron when presented a stimulus at the contrast level *c* ∈ {1,...,*C*} and size level *s* ∈ {1,...,*S*} as 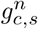, we compute the following suppression and contrast indices to numerically evaluate the tuning strength:

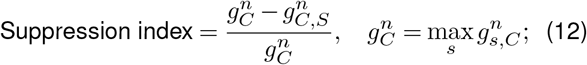

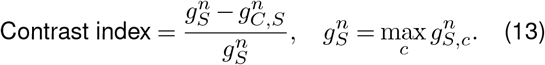

These two indices are highly correlated (*ρ* = 0.95, *p* < 0.001) which is why we use only the suppression index for the analysis. This correlation apparently stems from using Gabors as stimuli for this experiment. Indeed, increasing the size of a Gabor also increases the range of each pixel value, which is a similar effect to increasing the contrast.

#### Plaid stimuli

Another experiment we do with optimal Gabor stimuli is the one aimed at probing neurons or clusters for a potential effect of cross-orientation inhibition. To do some we construct plaid stimuli by superimposing two Gabors on each other, the optimal one and the one orthogonal to the optimal one, while varying the contrasts of both of these stimuli. Examples of such stimuli are shown in Fig. 2. The corresponding tuning curves allow us to see potential non-linear suppressing effect of increasing contrast of the orthogonal stimulus known as cross-orientation inhibition.

Denoting predicted activity of *n*-th neuron when presented a plaid stimulus with an optimal Gabor at the contrast level *p* ∈ {1,...,*P*} and an orthogonal Gabor at the contrast level *o* ∈ {1,..., *O*} as 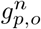, we compute the following numerical index to quantify the tuning strength:

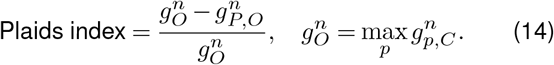

#### Comparison of optimal Gabor, DoG, and MEI stimuli

Optimal MEIs capture a wide variety of patterns in the receptive field, while optimal Gabors and DoGs are explicitly designed to represent a particular pattern. Comparing the responses to these stimuli allows us to quantify to what extent the receptive field captured by the MEI can be modelled by oriented gratings (Gabors) or center surround patterns (DoG).

For every neuron we normalize all three (Gabor, DoG, MEI) optimal stimuli to have same energy (*L*_2_ norm) at *E* different energy levels; such a normalization ensures that the stimuli have approximately the same contrast. Denoting predicted activity of *n*-th neuron when presented a Gabor, DoG or MEI at the energy level *e* as 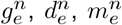 respectively, we compute the following summary statistics for comparing the responses to these stimuli:

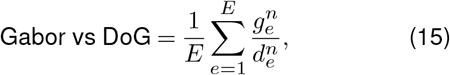

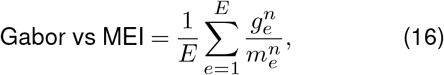

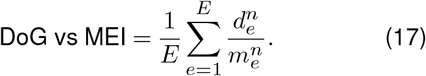

#### Non-linearity index

For every neuron we quantify the nonlinearity computations implemented by this neuron by comparing the predictions of the GLM and the CNN models. Specifically, for the *n*-th neuron we denote the GLM predictive correlation on the test set as 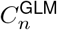, and the same quantity computed for the CNN model as 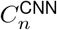. Then we compute the non-linearity index as follows:

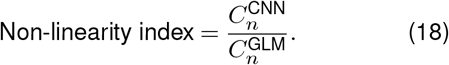

### In vivo verification

#### Experimental details

The experimental setting for the ImageNet scan in vivo verification was the same as in the main experiment (see above). The Gabor stimuli were presented in maximum contrast in 5 different sizes (*σ* ∈ [8,13.2,21.8,35.9,59.3]).

#### Selection of suboptimal stimuli

We find a small number of Gabor stimuli activating many neurons to a certain percentage of their optimal Gabor activities (Fig. 5D-E) using the following procedure:

- We compute predicted activities for every neuron for a large selection of Gabors located in the center of the image (the same set of stimuli that we used for finding the optimal Gabor apart from the differences in spatial locations; see above),
- We greedily choose the Gabor that activated most of the neurons to a given percentage of their optimal Gabor activity until we selected the required number of stimuli (we use 20 for the experiment).

## Data availability

All figures were generated from raw or processed data. The data generated and/or analyzed during the current study are available from the corresponding author upon request. No publicly available data was used in this study. All code and data will be available online upon the publication.

## Code availability

Experiments and analyses were performed using custom software developed using the following tools: ScanImage 2018a (Pologruto et al., 2003), CaImAn v.1.0 (Giovannucci et al., 2019), DataJoint v.0.11.0 (Yatsenko et al., 2015, 2018), TensorFlow v.1.15.0 (Abadi et al., 2015), NumPy v.1.17.3 (Van Der Walt et al., 2011), SciPy v.1.5.4 (Virtanen et al., 2020), Docker v.19.03.12 (Merkel et al., 2014), Matplotlib v.3.1.1 (Hunter, 2007), seaborn v.0.11.1 (Waskom, 2021), pandas v.1.1.5 (McKinney et al., 2010) and Jupyter Notebook v 6.0.1 (Kluyver et al., 2016). The code will be publicly available upon the publication.

## Animal research statement

All experimental procedures complied with guidelines approved by the Baylor College of Medicine Institutional Animal Care and Use Committee (IACUC).

